# Protracted morphine withdrawal corresponds with sex-specific alterations to motivated behavior and mesoaccumbal subcircuit dopamine cell plasticity

**DOI:** 10.1101/2023.01.29.526129

**Authors:** Devan M. Gomez, Taytum Kahl, Emily Berrington, Matthew C. Hearing

## Abstract

**Background:** Opioid use disorder is associated with enduring psychological withdrawal symptoms believed to contribute to drug abuse. Amongst these are shifts in motivational states, wherein pursuit of drug consumption exceeds that of non-drug rewards, reinforcing escalated opioid use and relapse vulnerability. A critical regulator of behavioral reinforcement, the mesoaccumbal dopamine (DA) system is thought to be both necessary and sufficient for opioid motivation. However, previous research into its involvement in opioid withdrawal has been limited to acute vs protracted timepoints, global neuroadaptations vs those in subcircuits, and overwhelmingly focused on males vs females.

**Methods:** Evaluations of effort-based motivated behavior for both sucrose and morphine reward were combined with patch clamp electrophysiological assessments of synaptic plasticity within lateral vs medial DA neurons projecting to the lateral vs medial nucleus accumbens shell during protracted morphine withdrawal in male and female mice. Further effects of mesoaccumbal subcircuit inhibition on motivated behavior for sucrose were also measured.

**Results:** Protracted morphine withdrawal was found to be associated with elevations in morphine seeking, intake, and motivation compared to saline controls in both sexes. Escalation of intake was paralleled by a male-exclusive reduction in motivation for the non-drug reward, sucrose. Male-exclusive neuroadaptations during protracted withdrawal were also found, with reductions in neuronal excitability and increased inhibitory (GABA_A_R-dependent) synaptic transmission found in lateral ventral tegmental area (VTA) DA neurons projecting to the lateral nucleus accumbens shell, though not in medial DA projections to the medial shell. Finally, chemogenetic inhibition of the lateral but not medial subcircuit was found to significantly reduce motivated responding for sucrose in male morphine-naïve mice.

**Conclusions:** These data suggest that protracted opioid withdrawal is associated with a sex-independent increase in opioid consumption and motivation. They also suggest that male-specific reductions in motivation for non-drug reward during protracted withdrawal may be driven by a hypoactive state in a lateral mesoaccumbal DA subcircuit driven in part by increased inhibition of DA cells. These insights may be useful in development of therapies that temper withdrawal-associated psychological states predisposed towards prolonged and escalated opioid intake, a major treatment goal for OUD patients.

## INTRODUCTION

Opioid abuse poses a serious health threat in the US as rates of opioid use disorder (OUD) and related overdoses continue to rise. A chronic disease, OUD afflicts patients with enduring relapse vulnerability at least partially through adverse withdrawal symptoms. Clinical and preclinical findings alike indicate prolonged, or protracted withdrawal, encompasses a set of negative affective symptoms including dysphoria, anhedonia, and altered reward processing ^1,2^. Unlike the more widely studied acute somatic withdrawal symptoms, which subside in a matter of days, protracted withdrawal symptoms may endure for months to years to impart a long-lasting susceptibility to relapse and continued drug intake ^3^.

Altered reward processing and motivation is a key symptom of protracted withdrawal that is described by the Diagnostic and Statistical Manual of Mental Disorders (DSM-5) as diagnostic criteria for OUD ^4^. In particular, the DSM-5 describes a type of motivational switch during OUD, through which patients turn their efforts away from obtaining non-drug rewards towards opioids themselves. As such, this type of affective symptomology may strongly indicate a patient’s likelihood to abuse opioids. Yet, to the authors’ knowledge, no investigation has been focused on comparing an organism’s effortful motivation for natural and drug reward during protracted opioid withdrawal.

One of the brain’s central reward centers is the dopamine (DA) mesoaccumbal system, containing DAergic projections from the ventral tegmental area (VTA) to the nucleus accumbens (NAc). Subserving reinforcement across species, mesoaccumbal DA activity has been widely implicated in reward processing and motivated behavior ^5–10^. This system is also implicated in regulating aversion and negative affect associated with drug-seeking behavior and withdrawal ^2,11–14^. Increasing evidence suggests these behavioral processes are driven by anatomically and functionally distinct populations of DA neurons within the mesoaccumbal system. And recent findings have identified a causal link between activation of DA cells selectively projecting to the medial NAc shell and opioid reinforcement ^15^. While opioid-induced changes to motivational circuitry have been researched in the context of acute reward and withdrawal, little attention has been focused on protracted withdrawal and the neuroadaptive alterations underlying lasting changes to mood and behavior.

## MATERIALS & METHODS

### Animals

Adult male and female wild-type C57BL/6 mice were bred in-house or commercially purchased (Jackson Laboratory) and maintained in a temperature and humidity-controlled room. For behavioral studies, male and female wild-type mice (postnatal day 52-72 at beginning of training) were used. For electrophysiology, adult male and female wild-type (postnatal day 68-114 at time of recording) were bred in-house with Pitx3-EGFP transgenic mice, generously provided by Dr. Kevin Wickman. Only heterozygous mice were used for experiments. For chemogenetic studies, DAT-IRES-Cre mice (Jackson Labs) were bred in-house with wild-types mice and only heterozygous mice were used (postnatal day 52-85 at beginning of training). All mice were single housed for all experiments on a 12-hour light/dark cycle with food and water available ad libitum, with all experiments run during the light portion. All experiments were approved Institutional Animal Care and Use Committee at Marquette University.

### Escalating morphine injections

To induce a physiologically dependent state within both male and female mice, an escalating morphine regime was used as previously described ^16^. Across a period of 5 days, 2 injections (saline or morphine) were given each day in their home cage approximately 12 hours apart, with doses for each day at 20, 40, 60, 80, and 100 mg/kg (10 injections total).

### Spontaneous somatic withdrawal scores

Scoring of acute somatic withdrawal scores were taken as previously described ^16^. Twenty-four hours and 14 days following the final injection, mice were placed into a clear plastic cage (16.5”×8”×7.5”) while signs of somatic withdrawal were hand scored during a 30 min period. Somatic measures were chosen based on previous works examining morphine withdrawal ^17–19^. Jumps, tremors, paw flutters, wet dog shakes, piloerection, and grooming were hand scored with each measurement recorded as a score of one, including the singular possible observation of piloerection, to generate a global withdrawal score with the following equation: *jumps3+tremors+grooming+pawflutters+shakes+pilerection*. Scorer was blind to treatment when possible, occurring for approximately half of all scores.

### Sucrose preference

Mice were habituated to single housing in cages with constant access to two drinking bottles, both containing water. Following 5 days of habituation, one bottle was filled with 1% sucrose solution and each bottle was weighed prior to exposure to preference testing. This concentration was chosen as it has been previously used in sucrose preference tests of anhedonia ^20,21^. Sucrose and water bottle placement was alternated from the left and right side between cages. Exposure to both bottles lasted for 24 days, after which each bottle was weighed to measure consumption. This measurement was taken for each mouse before morphine injections were given (baseline) and again 11 days following the final injection of either an escalated morphine regimen or saline (withdrawal).

### Stereotaxic surgeries

All surgical procedures were performed with mice under general isoflurane anesthesia (1-3%) prior to the subject being placed in the stereotaxic frame (Kopf Instruments, Tujunga, CA, USA). For intraccumbal injection of both Retrobead™ and DREADDs, injections were targeted to either the medial (bregma coordinates: anterior/posterior, + 1.45 mm; medial/lateral, + 0.55 mm; dorsal/ventral, −4.4 mm) or lateral (anterior/posterior, +1.45 mm; medial/lateral +/− 1.75 mm; dorsal/ventral −4.9 mm) nucleus accumebns shell. Both red and green colored Retrobeads™ were injected in the majority of Pitx3-EGFP animals used for electrophysiology studjes, with different colors alternately assigned to either the medial or lateral nucleus accumbens shell. A subset were injected with only one color in either or both subregions.For GiDREADD studies, either AAV5-hM4Di-hsyn-DIO-mCherry or AAV5-hsyn-DIO-EYFP (Addgene) was injected targeted in DAT-IRES-Cre mice in either the medial or lateral nucleus accumbens shell using the same coordinates as above. This approach leveraged the retrograde tendencies of the AAV5 serotype in a Cre-inducible virus, enabling pathway-selective targeting of dopamine cells in DAT-IRES-Cre mice. Mice were allowed a minimum 5-day recovery period before beginning behavior testing.

### Intravenous catheterization surgeries

Mice were implanted with an intravenous standard mouse jugular vein catheter (Access Technologies; 2/3Fr. x 6cm silicone, Cat No. AT-MJVC-2914A) connected to a backmount (PlasticsOne; 22GA, Part No. 8I31300BM01) under general isoflurane (1-3%) anesthesia. All mice underwent surgery approximately 5 days following their last injection of either morphine or saline vehicle. Following catheter surgery, mice were single-housed and allowed to recover for at least 5 days prior to beginning self-administration. During this time, mice were acclimated to handling and catheter manipulation. For all self-administration sessions, catheters were flushed with 0.05 mL (i.v.) of heparinized (20 IU/ml; Hospira, Inc.) bacteriostatic 0.9% saline (Hospira, Inc.) containing gentamicin sulphate (Sparhawk Laboratories, Inc.; 0.25 mg/ml) immediately before and containing enrofloxacin (Norbrook Laboratories; 4.994mg/mL) immediately after each session.

### Sucrose self-administration

Operant training occurred in standard conditioning chambers fitted with levers and cue lights, housed in sound-attenuating compartments (Med Associates). Two levers flanked a food magazine, wherein 20% liquid sucrose reward was delivered after each appropriate response. A cue light positioned directly above the active lever was illuminated and the house light turned off for 20 seconds after each press during training sessions, which were under a fixed ratio 1 schedule of reinforcement. During FR1 training, each active lever (left lever) press was reinforced with reward, while each inactive lever (right lever) press resulted in neither cue light illumination nor reward delivery. Following reward delivery, a 20 second timeout period was initiated, during which active lever pressing resulted in no reward until the period ended. Sessions lasted for 60 minutes, after which all manipulanda were terminated and the animal was shortly removed.

Animals were initially trained under food deprivation (90-85% their free-feeding weight) daily until they responded with ≥ 25 active presses for 3 consecutive days. Animals were then returned to ad libitum feeding and continued FR1 training for 10 consecutive days. Low responding animals after this point (<25 presses) were removed from the study. Twenty-four hours following completion of this training protocol, animals underwent progressive ratio tests for 20% liquid sucrose reinforcer as baseline motivational assessments of natural reward. During progressive ratio tests, an exponentially increasing schedule of reinforcement (5e.2*n)−5) was used, wherein each successive reward required a higher response ^22^. Testing continued until 30 minutes elapsed with no response or a total of 180 min passed. Standardization of sucrose breakpoint difference scores: using saline control values as a standard, the average difference between each individual score within the morphine group and the mean of the corresponding saline group (same timepoint and sex) was calculated.

### Morphine intravenous self-administration

Each animal was subjected to escalated morphine or saline, as described above, 3 – 24 hours following their initial (baseline) sucrose progressive ratio test. Following a home cage forced abstinence period of 11 days, mice ran a second post-treatment sucrose progressive ratio test. All animals were then assessed for intravenous catheter patency using .05 mL of either 5 or 9 mg/kg sodium brevital solution through their catheter line. Non-patent animals were identified by lack of immediate sedation and removed from further testing. Patent mice started 2-hour intravenous morphine self-administration sessions (0.025mL of 0.1mg/kg/infusion) in the same operant chambers used for sucrose 24-hours following their last sucrose progressive ratio test. Daily 2-hour morphine sessions were run for 10 days. Progressive ratio tests were run for morphine identical to sucrose on the 3^rd^ day of self-administration and 24-hours following the last 2-hour session (11^th^ and final day). This timeline allowed animals to acquire morphine intravenous self-administration and adjust to the change in reinforcer before effortful motivation for morphine was measured.

### Chemogenetic manipulation of sucrose self-administration

Following intra-cranial surgeries, operant training for 20% liquid sucrose took place as above. Animals were initially trained under food deprivation (90-85% their free-feeding weight) daily until they responded with ≥ 25 active presses for 3 consecutive days. Animals were then returned their standard chow and continued FR1 training for 10 consecutive days fed ad libitum. Low responding animals after this point (<25 presses) were removed from the study. Twenty-four hours following completion of this training protocol, animals underwent progressive ratio tests for 20% liquid sucrose reinforcer 30 minutes after intraperitoneal injection of saline vehicle. Twenty-four hours following this, animals were given a second progressive ratio test 30 minutes after intraperitoneal injection of CNO (2 mg/kg). During progressive ratio tests, an exponentially increasing schedule of reinforcement ((5e.2*n)−5) was used, wherein each successive reward required a higher response (Richardson & Roberts, 1996). Testing continued until 30 minutes elapsed with no response or a total of 180 min passed.

### Mobility Assessments

At least twenty-four hours following the final injection, mice were placed into a clear plastic cage (16.5”×8”×7.5”) in a dimly lit room and measured for open field mobility during 20 minute sessions. Cages were separated with opaque plastic dividers to ensure mice were unable to visually detect each other when more than one animal was run at a time. Two daily 45-minute habituation sessions preceded each test for each animal. Twenty-four hours following habituation, an initial baseline test was run 30-minutes after saline injection (IP), followed by a second and final test 24-hours later (30-minutes after 2 mg/ kg CNO IP injection). Cameras placed above the cages recorded each animal’s movement while ANY-maze software (Stoelting) calculated average speed (m/s), total distance (m) and total rotations of each animal.

### Slice electrophysiology

Electrophysiological measurements of VTA dopamine neurons occurred in acute slice preparations from behaviorally-naive adult Pitx3-EGFP mice (68–114 d) receiving an intra-NAc infusion of Red or Green Retrobeads^®^. Horizontal slices (225mm) containing the VTA were prepared in the following ice-cold high sucrose substituted ACSF: 220 mM Sucrose, 2.5 mM KCl, 1.25 mM NaH2PO4, 25 mM NaHCO3, 11mM glucose, 2 mM MgCl2, and 2 mM CaCl2 (pH 7.4, 300 mOsm). Slices were allowed to recover at room temperature in the following ACSF for at least 45 minutes prior to recordings: 125 mM NaCl, 2.5 mM KCl, 1.25 mM NaH2PO4, 25 mM NaHCO3, 11mM glucose, 1 mM MgCl2, and 2 mM CaCl2 (pH 7.4, 300 mOsm). Acute brain slices were gravity perfused with oxygenated ACSF at a temperature of 29°C-33°C using at a flow rate of ∼2-2.5 ml/minute. Neurons in either the medial or lateral VTA exhibiting appropriate dual fluorescence were targeted for recordings. Sutter Integrated Patch Amplifier (IPA) with Igor Pro (Wave Metrics, Inc.) was used for the data acquisition software. Recordings were filtered at 2kHz and sampled at 5kHz for current-clamp recordings and voltage-clamp recordings. Miniature postsynaptic current recordings were filtered at 2kHz and sampled at 20kHz. IPSCs/EPSCs were analyzed using MiniAnalysis software using a 5pA detection threshold (Synaptosoft). Rheobase was assessed using a 20pA 1-second current-step injection from -120-600pA. Rheobase was defined as the minimum current step that evoked at least one action potential. Spontaneous EPSCs and IPSCs were assessed at a holding potential of -72mV. For the purposes of EPSCs, this is past the calculated ECl- of - 58mV, enabling isolation of AMPA-receptor mediated currents without the need of GABA_A_-receptor antagonists. Series and input resistances were tracked throughout the experiment. If series resistance increased >20% or reached 40MΩ, the experiment was excluded from analysis. Recordings were filtered at 2kHz and sampled at 5kHz for current-clamp recordings and voltage-clamp recordings. Miniature postsynaptic current recordings were filtered at 2kHz and sampled at 20kHz. Rheobase recordings utilized borosilicate glass pipettes filled with an internal solution of potassium gluconate as previously described: 140mM K-Gluconate, 5.0mM HEPES, 1.1mM EGTA, 2.0mM MgCl2, 2.0mM Na2-ATP, 0.3mM Na-GTP, and 5.0mM phosphocreatine (pH 7.3, 290mOsm) ^23,24^. For mEPSCs, borosilicate glass pipettes were filled with cesium methosulfate internal solution as previously described: 120mM CsMeSO4, 15mM CsCl, 10mM TEA-Cl, 8mM NaCl, 10mM HEPES, 5mM EGTA, 0.1mM spermine, 5mM QX-314, 4mM ATP-Mg, and 0.3mM GTP-Na ^23,24^. For mIPSCs, borosilicate glass pipettes were filled with calcium choride internal solution: 120 mM CsCl, 10 mM HEPES, 2 mM MgCl2, 1 mM EGTA, 2 mM NaATP, 0.3 mM NaGTP, 1 mM QX-314 (pH 7.4, 294 mOsm). Miniature IPSCs were recording with addition of 10µM NBQX to the bath.

### Histological assessment of Retrobead™ & virus expression

Accuracy and selectivity of Retrobead and virus targeting was assessed in slices prior to electrophysiology studies or post hoc in fixed tissue. Assessment of Retrobead injection placement was verified before electrophysiology took place for each animal either by eye, utilizing a flash equipped camera, or under a bright-field microscope when necessary. Post hoc viral placement was verified by fixing brains in 4% paraformaldehyde for 24 hours following transcardial perfusion with 4% paraformaldehyde, then placed in phosphate buffered saline until 50µm slices were obtained using a vibratome. Slices were then washed 7×3 minutes in 0.05M PBS, incubated in 0.05M PBS with .3% Trition X-100 for 1 hour, and incubated in the following blocking solution for 3 hours at room temperature: 1% BSA, 5% Normal Goat Serum, 0.3M Glycine, and .1% Triton X-100. Incubation with primary antibody for 36 hours at 4°C then took place in the following solution: 1% BSA, 5% Normal Goat Serum, .1% Triton X-100, 1:300 anti-tyrosine hydroxylase antibody (EMD Milliipore), and 1:2000 anti-mCherry antibody (abcam). Slices were then washed 6×10 minutes in .05M PBS and incubated with both Alexa-Fluor 488 and 555 secondary antibody (1:1000; Thermo Fisher Scientific) in .05M PBS with .1% Triton x100 for 2 hours at room temperature. Final 5×5 minutes washes followed in .05M PBS before sections were mounted on glass cover slips and applied with ProLong Diamond Antifade Mountant with DAPI (Thermo Fisher Scientific). Slides were allowed to dry for 24-hours before being imaged using a Nikon Eclipse Ti2 confocal microscope. Only mice with virus expressed bilaterally in the appropriate VTA subregion were used in analyses.

### Statistical Analysis

Statistics for t-tests were performed using GraphPad Prism 8.4. ANOVA analyses were performed using SPSS, with repeated measures on day. Bonferroni post hoc comparisons were conducted when necessary. The threshold for statistical significance was p<0.05. Data are expressed as mean ± SEM. Partial eta squared values are denoted by η_**p**_^**2**^.

## RESULTS

### Dependence-indicating morphine withdrawal signs dissipate after 14 days of forced abstinence

Opioid dependence is identified by characteristic somatic withdrawal signs following acute abstinence ^16,25,26^. To verify induction of morphine dependence in both male and female mice, spontaneous somatic withdrawal signs were scored both 24-h and 14-d after the last injection of either morphine or saline vehicle (**Figure 1A**). Morphine treated male and female mice displayed significant increases in acute somatic withdrawal signs compared to their saline counterparts, with a 2-way repeated measures (RM) ANOVA revealing a significant interaction between day and treatment (F(1,57)=24.373, p<.001, η_**p**_^**2**^=.300). Post hoc analysis revealed treatment effects at 24-h for both males (F(1,57)=16.883, p<.001, η_**p**_^**2**^=.229) and females (F(1,57)=4.293, p=.043, η_**p**_^**2**^=.070), as well as significant decreases in this effect during protracted withdrawal (males: F(1,57)=53.120, p<.001, η_**p**_^**2**^=.482; females: (F(1,57)=193485, p<.001, η_**p**_^**2**^=.255), indicating that significant morphine-induced increases in withdrawal symptoms across sex comparably diminished during protracted withdrawal (**Figure 1B**).

**Figure 1.**
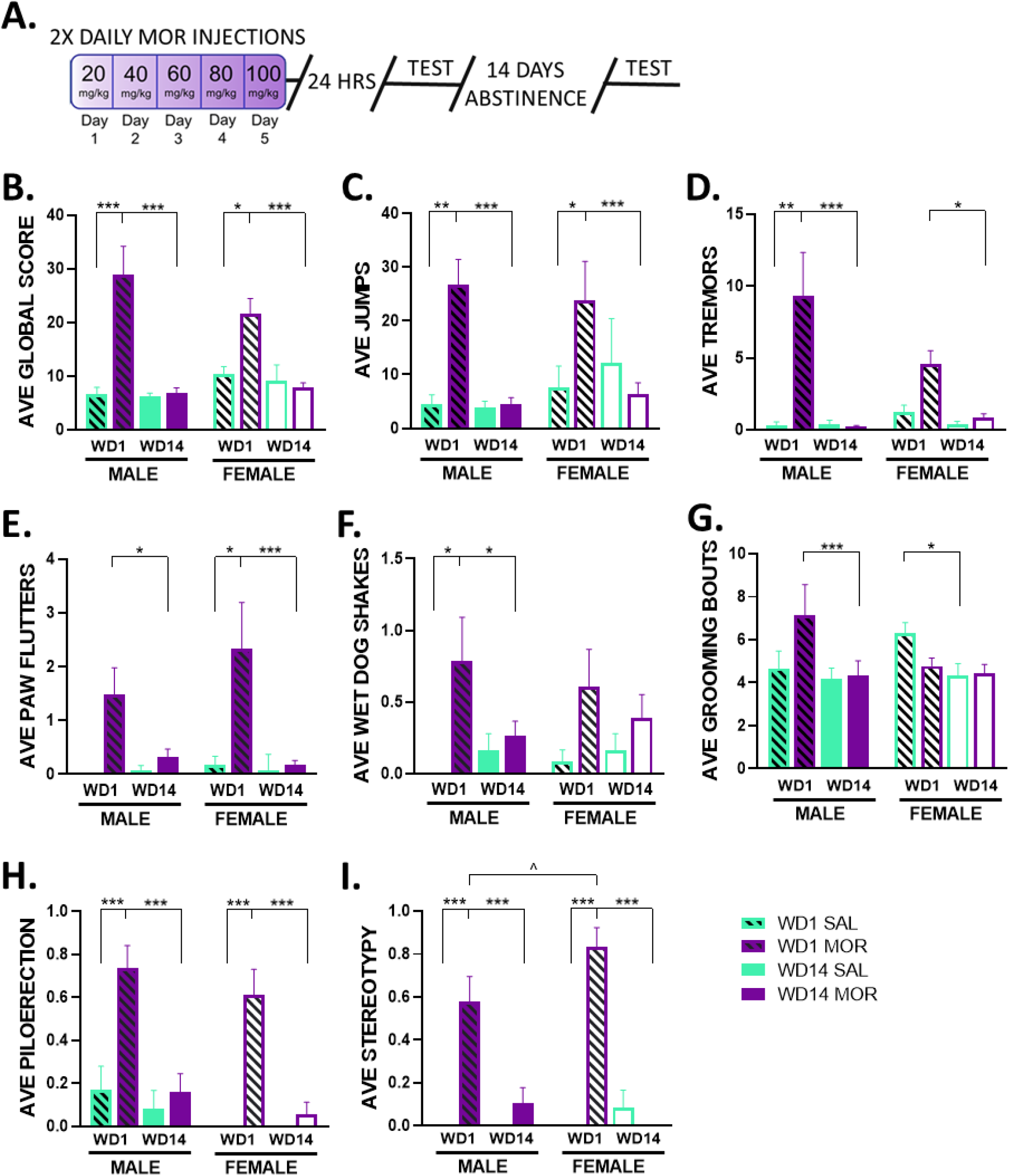
Acute somatic withdrawal signs fade during protracted morphine withdrawal. **(A.)** Schematic of experimental timeline including an escalating morphine injection regimen (2X daily injections, 5 days, s.c.) and assessment of withdrawal symptoms 24 hours (filled bars; left) and 14 days (open bars; right) following the final injection. **(B.)** Global somatic withdrawal signs assessed 24-h or 14-d post injections in in male (left; filled) and female (right; unfilled) treated with morphine (purple) or saline (green). Individual withdrawal signs of **(C.)** jumps, **(D.)** tremors, **(E.)** paw flutters, **(F.)** wet dog shakes, **(G.)** Grooming bouts **(H.)** piloerection, and **(I.)** stereotypy 24-h and 14-d following injections. N=12-19 per group. *p< 0.05. **p< 0.01. ***p< 0.001

Alternatively, post hoc comparisons revealed no marginal effects of sex (F(1,57)=1.202, p=.278) or treatment (F(1,57)=1.725, p=.194) on day 14 of withdrawal, indicating that morphine-induced somatic withdrawal sings were indeed visibly absent during protracted withdrawal (**Figure 1B**).

Analysis of the impacted of individual withdrawal symptoms showed a similar dissipation of significant sex-independent, morphine-related acute symptomology, including day x treatment interactions for jumps (F(1,57)=19.711, p<.001, η_**p**_^**2**^=.257), tremors (F(1,57)=9.281, p=.004, η_**p**_^**2**^=.140), paw flutters (F(1,57)=8.589, p=.005, η_**p**_^**2**^=.131), and piloerection (F(1,57)=19.107, p<.001, η_**p**_^**2**^=.251) (**Figure 1C, D, E & H**). The only individual withdrawal signs for which sex interacted with treatment effects were grooming bouts and instances of stereotypy, each showing a day X treatment X sex interaction [grooming: (F(1,57)=5.163, p=.027, η_**p**_^**2**^=.251); stereotypy: (F(1,57)=4.008, p=.050, η_**p**_^**2**^=.066)]. Post hoc tests revealed a trend of larger mean grooming bouts in male (F(1,57)=3.541, p=.065, η_**p**_^**2**^=.058) and significantly larger mean strereotypy value in female (F(1,57)=4.781, p=.033, η_**p**_^**2**^=.177) withdrawn mice during 24-h withdrawal only (**Figure 1G,I**). Furthermore, only male withdrawn mice showed significant decreases in mean grooming bouts across tests days (F(1,57)=13.552, p<.001, η_**p**_^**2**^=.192) while female saline controls displayed decreased values across days (F(1,57)=4.238, p=.044, η_**p**_^**2**^=.069) (**Figure 1G**). Taken together, these data indicate the escalating dose regimen of morphine promotes significant somatic withdrawal that can be observed 24-h following treatment and subsequently dissipates with more protracted withdrawal.

To assess changes in weight induced by their morphine treatment across withdrawal, weights were tracked daily for both sexes across the injection regimen and 14-day withdrawal period. A 2-way RM ANOVA revealed significant interactions between day and treatment (F(18,17)=15.575, p<.001, η_**p**_^**2**^=.943) as well as sex, day and treatment (F(18,17)=2.723, p=0.022, η_**p**_^**2**^=.742). Post hoc comparisons found that significant morphine-associated decreases in weight were restricted to treatment days and acute withdrawal, with weight rebounding to levels at or near saline counterparts soon after. Specifically, morphine treated males showed differences on injection days 3-5 as well as the first 2 days of withdrawal. Morphine treated females showed a similar pattern with exception to a quicker rebound in weight, with only the first day of withdrawal accompanying a significant decrease (**Figure 2**). These results suggest morphine dependence-induced weight loss is reversed during protracted withdrawal in both sexes.

**Figure 2.**
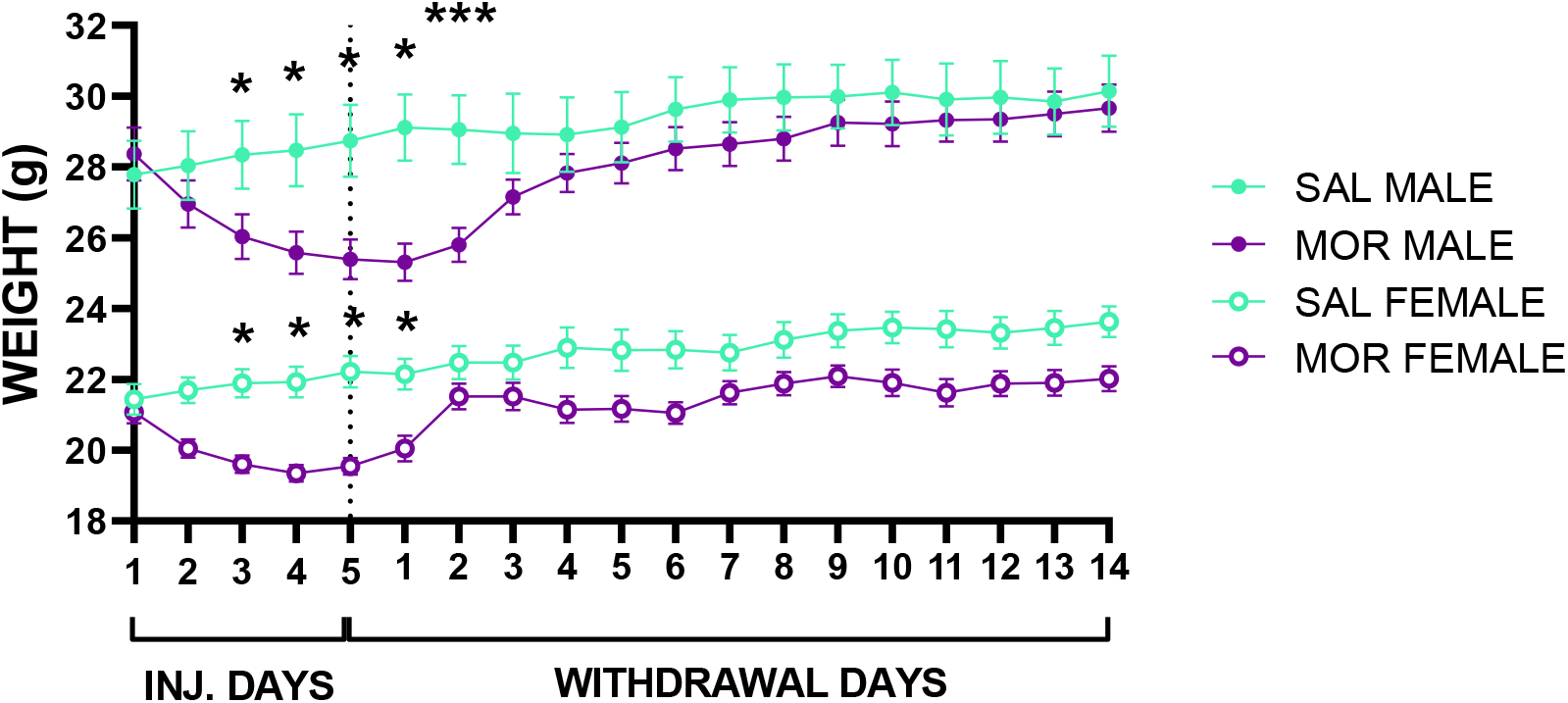
Morphine-induced decreases in body weight rebounded similarly for both sexes following protracted withdrawal. Morphine (purple) but not saline (green) treated male (solid dots) and female (open dots) mice significantly lost weight during the last 3 days of injections and the first to second day of acute withdrawal. Weights for moprhine treated mice rebounded for both sexes after this point, reaching levels similar to their respective saline counterparts. N=11–16 per group. *p< 0.05, ***p< 0.005 comparing within-sex body weights on specific days.

### Protracted morphine withdrawal corresponds with sex-specific decreases in effortful responding for sucrose

Protracted morphine withdrawal has been shown to alter effortful motivation for sucrose reward in rats, although reports have been mixed and focused selectively in males ^26–29^. To assess withdrawal-associated motivational changes for sucrose in male and female mice, progressive ratio (PR) tests were run in animals before morphine or saline treatment to assess baseline motivation within each animal, and again during protracted withdrawal (**Figure 3A**). Initial analysis of the within-subject breakpoint values revealed considerable variability, particularly for females (**Figure 3B, C**). Breakpoints of the morphine-withdrawal groups were standardized relative to their sex-matched saline counterparts to illustrate the effect of morphine withdrawal. In males, this showed a reduction in breakpoint value across testing days in morphine withdrawn mice (−9.78). In contrast, female withdrawn mice displayed higher breakpoint values at both testing days, displaying a negligible increase following withdrawal (+0.42) (**Figure 3D**). When run for both sexes, RM ANOVAs of the standardized breakpoint data resulted in no interaction effect for either males (F1, 45)=2.161, p=.149, η_**p**_^**2**^=.046) or females (F(1, 36)=.004, p=.952, η_**p**_^**2**^=.000). However, post hoc comparisons did find a significant difference across test days in morphine treated male (F(1, 45)=4.232, p=.046, partial eta^2^=.086) but not female (F(1,36)=.007, p=.932, partial eta^2^=.000) mice, indicating the simple effect of day was significant for morphine male mice only. Furthermore, a t-test with Welch’s correction found a significant decrease in raw lever presses in male (t(42.7)=2.192, p>0.0340, d=-.637) but not in female (t(34.4)=1.388, p=.1740; d=.45) withdrawn mice (**Figure 3E**). These data indicate that while dependence was induced in both male and females, effortful responding for sucrose decreased in withdrawn males but not females compared to their saline counterparts.

**Figure 3.**
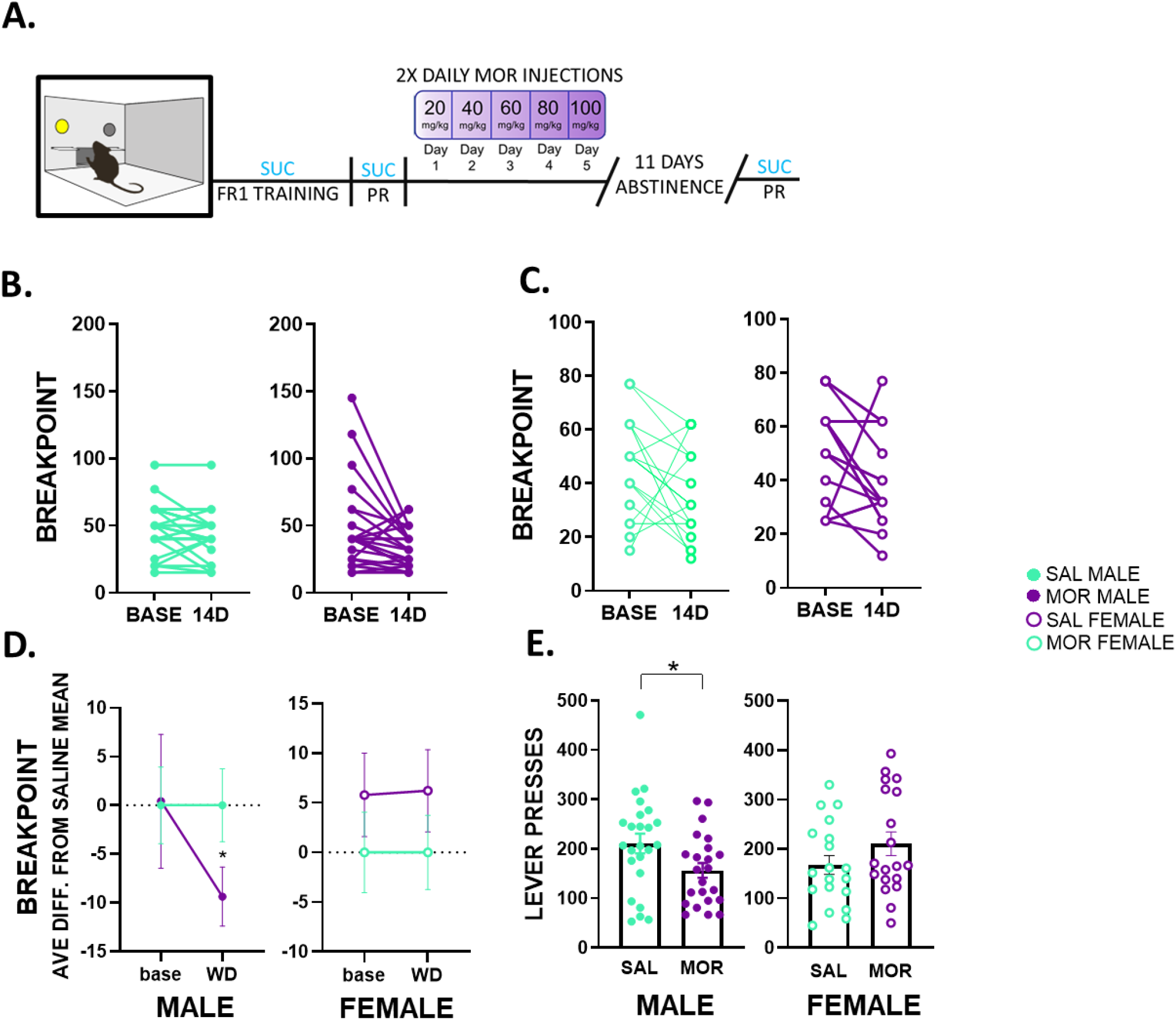
Males display decreased effortful responding for sucrose during protracted withdrawal. **(A.)** Male (filled dots) and female (empty dots) mice underwent progressive ratio tests for liquid sucrose both before morphine (purple) exposure and following 11 days of withdrawal or during same timepoints following saline (green) injections. **(B**., **C.)** Within-subject breakpoint data revealed high variability for females that obscured treatment effects. **(D.)** Standardized breakpoint values revealed a simple effect of day for male but not female withdrawn mice. **(E.)** Independent t-tests for raw lever presses revealed a significant decrease in male but not female withdrawn mice. N=19–24 per group. *p< 0.05.

### Hedonic behavior towards sucrose is unaffected by protracted morphine withdrawal

To investigate whether observed decreases in effortful sucrose responding reflect altered hedonics, a two-bottle sucrose preference test was run for a subset of animals. Both opioidergic and dopaminergic (DAergic) signaling within the mesoaccumbal system have been implicated in hedonic responses to sucrose reward ^30–32^. In rodents, anhedonia is typically identified by a reduction in the consumption of and preference for highly palatable rewards, such as sucrose, over natural food and/or water. Thus, we used a within-animal approach to measure sucrose versus water consumption as a measure of preference prior to morphine or saline injections and 14-d after treatment (**Figure 4A**). Two-way (sex x treatment) RM ANOVA revealed no significant effect of day (F(1,40)=1.078, p=.305), and no interactions between day and sex (F(1,40)=1.757, p=.193), day and treatment (F(1,40)=.002, p=.961), or day, treatment and sex (F(1,40)=.057, p=.813). This suggests protracted morphine withdrawal did not alter sucrose preference scores in either sex (**Figure 4B, C**).

**Figure 4.**
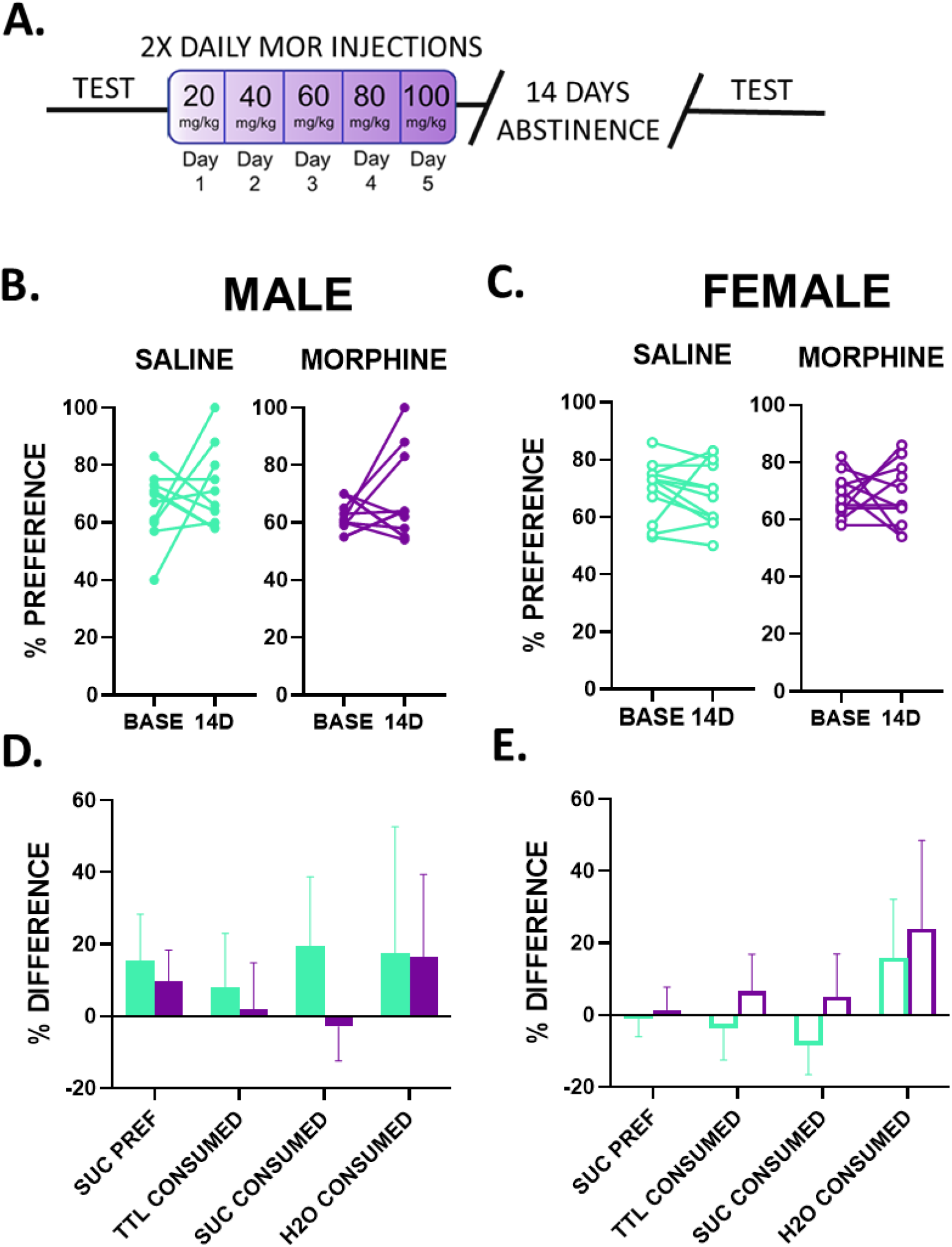
Protracted morphine withdrawal is not associated with altered hedonics. **(A.)** Male (filled dots/ bars) and female (empty dots/ bars) mice were tested for sucrose preference and consumption using a two-bottle choice test in their homecage both before morphine (purple) exposure and 14-days morphine withdrawal or similar timelines with saline vehicle (green). No significant difference in sucrose preference between pre-treatment baseline and protracted withdrawal days in were found in **(B.)** male or **(C.)** female withdrawn mice. Percent difference scores calculated for sucrose preference and total liquid, sucrose, and water consumed revealed no significant differences for either **(D.)** males or **(E.)** females. N=10–12 per group.

To clarify both the direction (increase vs decrease) and magnitude of change in either sucrose or water consumed following withdrawal, percent difference scores were calculated for both solutions using pre-treatment baseline measures as a standard. This allowed isolation of percent difference scores for sucrose, water, and total liquid consumed. Individual analysis of these scores using a 2-way ANOVA (treatment x sex) showed no significant difference in total liquid consumed, with no main effect of treatment or sex or interaction between them (**Figure 4D, E**). A difference in sucrose consumed between treatment groups was found in both males (saline: 19.629 ± 19.092, morphine: -2.893 ± 9.509, Hedges’ g=0.4747) and females (saline: - 8.403 ± 8.142, morphine: -5.022 ± 11.957, Hedges’ g=0.3932), though 2-way ANOVA analysis (treatment x sex) found no main effect of treatment (F(1,40)=.135, p=.715), sex (F(1,40)=.659, p=.422), or interaction (F(1,40)=2.105, p=.155) on percent difference scores (**Figure 4D, E**). These results suggest protracted morphine withdrawal does not significantly affect the hedonic value, or pleasurable effects, of sucrose or consummatory behaviors for this natural reward in a low-effort setting.

### Protracted morphine withdrawal increases intake and responding for morphine in both sexes

To determine whether prior dependence altered intake and motivation for morphine, mice were allowed to self-administer intravenous morphine (IVSA) 24-h after their final sucrose PR test. Under a fixed ratio schedule of reinforcement (FR1), mice underwent 2-hour sessions of morphine IVSA across 10 days, except for the 4^th^ day, during which an initial PR test for effort-based motivated responding was given (**Figure 5A**). Initial morphine PR tests were run on the 4^th^ day of IVSA to allow animals 3 days to adjust to the replacement of a sucrose reinforcer with intravenous morphine. Analysis of average infusions revealed highly skewed distributions across most session days. A logarithmic transformation of the data was chosen to adjust for this. Greenhouse-Geisser corrected results are reported as there was a violation of shericity. A 2-way (sex x treatment) RM ANOVA revealed a significant interaction between day and sex (F(4.015, 172.64)=2.693, p=.032, η_**p**_^**2**^=.059). Post hoc tests showed that this may be explained by high infusion rates of withdrawn males, with this group displaying significant withdrawal-related increases in mean infusions during 6 days of IVSA compared to trends in increased mean infusions for females during 3 days of IVSA (**Figure 5B**). Mean lever pressing data across days was also log transfomed as above. Greenhouse-Geisser corrected results are reported as adjustments to sphericity. Resulting 2-way RM ANOVA analysis showed a day x sex interaction (F(4.485, 192.854)=3.609, p=.005, η_**p**_^**2**^=.077). Post hoc tests showed significant withdrawal-related increases in mean lever presses during 5 IVSA days for males and 3 days for females, with males increasing their responding during the final days of IVSA and females displaying higher responding during the first 3 sessions (**Figure 5C**). These data suggest both sexes increase their morphine IVSA responding during protracted withdrawal.

**Figure 5.**
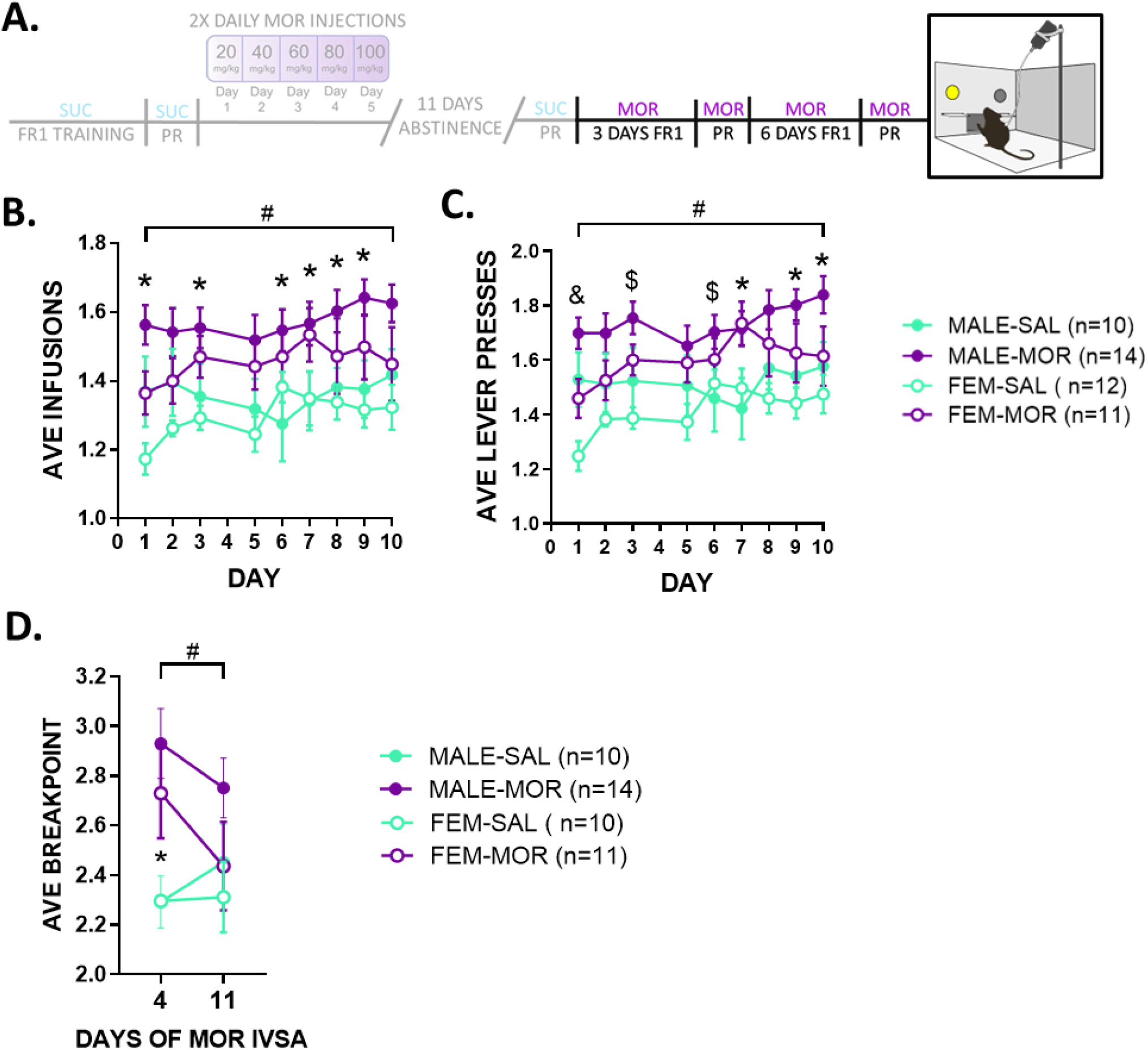
Protracted withdrawal increases intake, seeking & motivation for morphine across sexes. **(A.)** Male (solid dots) and female (empty dots) mice underwent daily 2-hour morphine (purple) or saline (green) IVSA sessions following sucrose assessments. Intake and total responding were measured daily while PR tests were taken during the 4^th^ and 11^th^ (final) day of IVSA. **<u>(B.)</u>** <u>Male</u> and female withdrawn mice displayed increases in average infusions across IVSA days, with males reaching significance across days 1, 3, 6, 7, 8 & 9 of IVSA. **(C.)** Significant increases in average active lever presses were found for both male (days 3, 6, 7, 9, & 10) and female (days 1, 3, & 6) withdrawn mice. **(D.)** Both sexes displayed significant increases in breakpoint values obtained for morphine reward during initial (4^th^ IVSA day) but not final (last IVSA day) PR tests as revealed by RM ANOVAs.. N=10-14 per group. *p< 0.05, **p< 0.005 comparing between-group on specific days. **#** indicates a main **(B**., **C.)** and marginal **(D.)** effect of day. **&** indicates a significant difference exlcusive to females. **$** indicates a significant difference shared by both sexes.

Assessments of morphine motivation during early self-administration (following 3-d IVSA) versus the 11^th^/ final day of IVSA were analyzed for both sexes using 2-way (sex x treatment) RM ANOVAs. To adjust for a skewed distribution, data was cube root transformed. Analysis revealed an interaction between day and treatment (F(1,39)=5.807, p=.021, η_**p**_^**2**^=.130). Post hoc tests showed significant increases in breakpoint values for withdrawn male (F(1,39)=11.726, p=.001, η_**p**_^**2**^=.231) and female (F(1,39)=5.026,p=.031, η_**p**_^**2**^=.114) mice during the first but not second PR test as well as a significant difference in values across days exclusively in female withdrawn mice (F(1,39)=4.165, p=.048, η_**p**_^**2**^=.096) (**Figure 5D**).

### Protracted morphine withdrawal is associated with male-exclusive neuroadaptations to mesoaccumbal subcircuit DA cells

Mesoaccumbal DA neurons are known to play a critical role in modulating reward, aversion, and reinforcement. Notably, while changes in DA cell physiology and synaptic regulation have been observed during acute opioid withdrawal, almost nothing is known regarding plasticity at more protracted time points. ^33–37,33^. As past studies assessing effects of morphine on DA neuron plasticity have used electrophysiological and neuropharmacological properties to identify DA neurons, our use of Pitx3-EGFP mice to genetically target VTA DA cells enabled excellent accuracy compared to much of the relevant literature ^34–37^. Following 14-d morphine withdrawal, we assessed changes in basal excitability (rheobase), excitatory postsynaptic currents regulated by AMPA-type glutamate receptors (sEPSCs), and inhibitory postsynaptic currents mediated by GABA-A type receptors(sIPSCs). Because intravenous morphine exposure increases DA release preferentially in the shell compared to the core subregion of the NAc ^38^, DA cells that project to the shell subregion were selectively targeted. Futher, due to evidence supporting distinct electrophysiological and behavioral reinforcement properties of VTA DA cell subpopulations determined by their NAc subregion projection bias ^6,39,40^, medial and lateral NAc shell-projecting cells were separately assessed using different color Retrobeads™ across both NAc subcompartments (**Figure 6A**).

**Figure 6.**
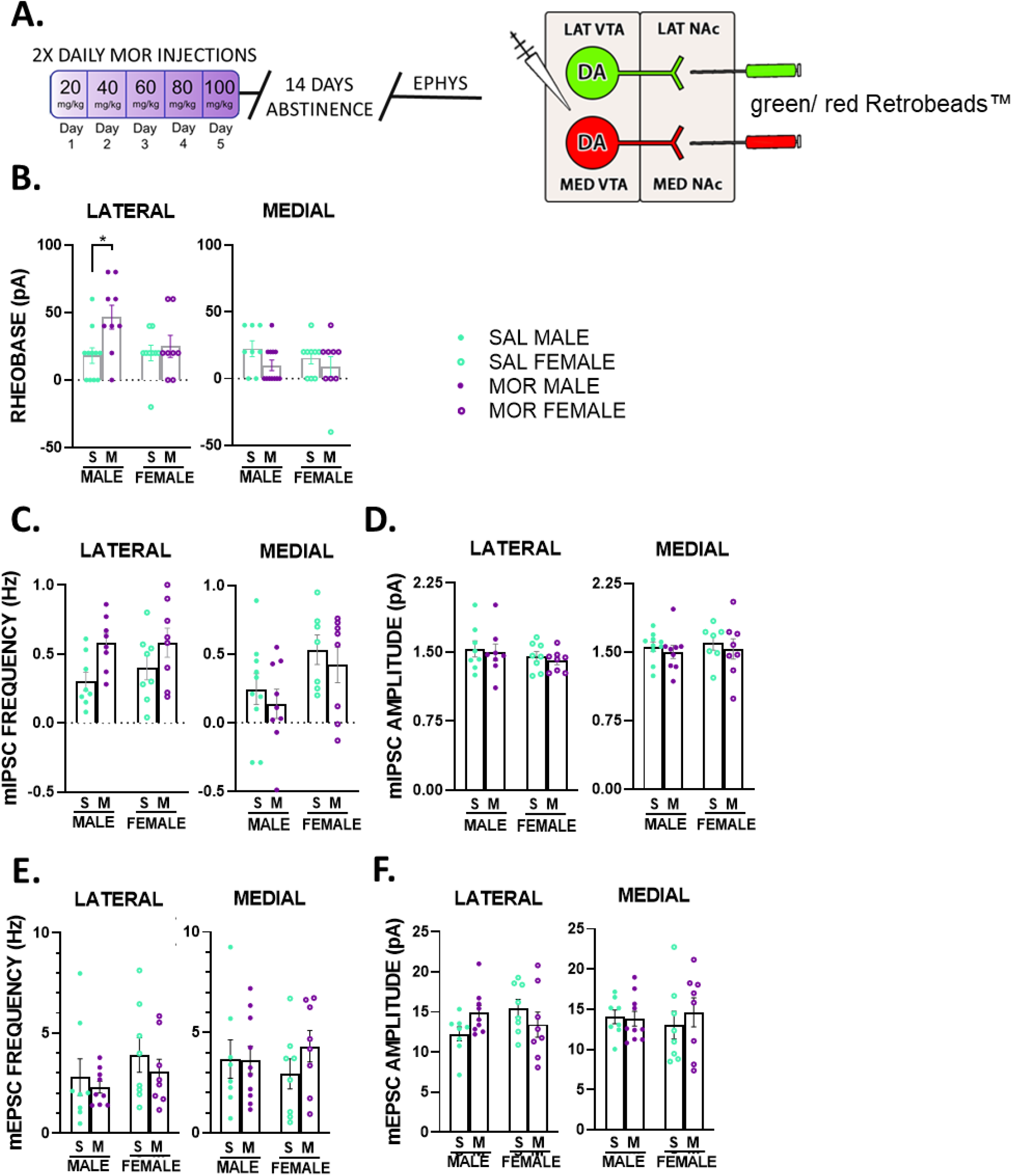
Male-exclusive neuroadaptations specific to lateral mesoaccumbal subcircuit VTA DA cells are associated with protracted withdrawal. **(A.)** Using different colored Retrobeads™, both lateral and medial NAc shell-projecting VTA DA cells were selectively targeted for whole cell recordings in male (solid dots) and female (empty dots) mice following 14-days morphine withdrawal (purple) or saline vehicle treatment (green). **(B.)** DA cells projecting to the lateral but not medial NAc shell (lateral subcircuit) displayed significantly increased rheobase in male withdrawn mice. **(C.)** Male withdrawn mice also displayed significant increases in mIPSC frequency selective to the lateral subcircuit. No significant treatment differences were found in **(D.)** mIPSC amplitude, **(E.)** mEPSC frequency or **(F.)** mEPSC amplitude in either sex. N=5–9 animals per group. n=8–12 cells per group. *p< 0.05.

Three-way (sex x subcircuit x treatment) ANOVA analysis of rheobase values showed a 3-way interaction (F(1,67)=4.962, p=.029, η_**p**_^**2**^=.069). Post hoc analysis revealed a significant treatment effect in the lateral VTA – to – lateral NAc shell (lVTA-lShell) subpopulation in males only ((F(1,67)=11.859, p<.001, η_**p**_^**2**^=.150). Further independent two-way ANOVAs for males and females showed significant interaction between treatment and subcircuit in males (F(1,36)=11.237, p=.002, η_**p**_^**2**^=.238), while females showed no interaction (F(1,31)=.054, p=817, η_**p**_^**2**^=.002), nor any main effect of treatment (F(1,31)=.368, p=.548, η_**p**_^**2**^=.012) or subcircuit (F(1,31)=.961, p=.334, η_**p**_^**2**^=.030). Bonferroni post hoc analysis of male rheobase data revealed a significant increase in withdrawn lVTA-lShell DA cells (F(1,36)=11.026, p=.002, η_**p**_^**2**^=.234) but not medial VTA – to – medial NAc shell (mVTA-mShell) DA cells (F(1,36)=2.059, p=.160, η_**p**_^**2**^=.054) (**Figure 6B**).

Initial analysis of sIPSCs revealed a highly skewed distribution, which was corrected by a logarithmic transformation of all the data. A resulting 3-way ANOVA run for sIPSC frequency detected an interaction between subcircuit and treatment (F(1,58)=5.599, p=.021, η_**p**_^**2**^=.088). Post hoc tests showed significantly higher mean frequency in lVTA-lShell compared to mVTA-mShell DA cells in male withdrawn slices only (F(1,58)=9.751, p=.003, η_**p**_^**2**^=.144) as well as a trend of treatment effect specific to male withdrawn lVTA-lShell DA cells (F(1,58)=3.771, p=.057, η_**p**_^**2**^=.061) (**Figure 6C**). Three-way ANOVA analysis of sIPSC amplitude data showed no significant interactions or main effects (**Figure 6D**).

Analysis of sEPSC frequency using a 3-way ANOVA revealed no interaction between sex, subcircuit and treatment (F(1,59)=.691, p=.409, η_**p**_^**2**^=.012), nor any main effect of sex (F(1,59)=.745, p=.392, η_**p**_^**2**^=.012), subcircuit (F(1,59)=1.388, p=.243, η_**p**_^**2**^=.023), or treatment (F(1,59)=.000, p=.999, η_**p**_^**2**^=.000) (**Figure 6E**), Similar analysis of sEPSC amplitude showed a trend in a 3-way interaction (F(1,59)=3.408, p=.070, η_**p**_^**2**^=.055) that post hoc tests indicated may best be explained by a trend in increased baseline amplitude in lVTA-lShell DA cells specific to females (F(1,59)=3.207, p=.078, η_**p**_^**2**^=.052). No main effects of sex (F(1,59)=.181, p=.672, η_**p**_^**2**^=.003), subcircuit (F(1,59)=.027, p=.870, η_**p**_^**2**^=.000) or treatment (F(1,59)=.315, p=.577, η_**p**_^**2**^=.005) were found (**Figure 6F**).

### Selective inhibition of lateral NAc shell-projecting DA cells decreases reinforcement of natural reward

The previously observed withdrawal effects in males, decreases in effortful responding for sucrose found concurrently with a hypoactive neural state, suggest withdrawal-related neuroadaptations may be causing the behavioral changes. Given the importance of DA to reinforcement behavior, it would be expected that inhibition of mesoaccumbal DA activity would decrease effortful responding for reward. But to the knowledge of the authors, no studies have focused on differences in effortful responding for sucrose following inhibition of different mesoaccumbal subcircuits. To pursue replicating the effect of decreased motivated behavior for sucrose during protracted withdrawal, a loss-of-function study was used in male morphine-naïve mice between mesoaccumbal subcircuits. For this, DAT-IRES-Cre mice received viral infusion of an inhibitory DREADD (AAV5-hsyn-DIO-hM4Di-mCherry) in either the lateral or medial NAc shell (**Figure 7A**). A Cre-dependent AAV-5 serotype viral construct was used for intra-accumbal infusions due to its tendency to transport retrogradely (Samaranch et al., 2017; Simmons et al., 2019). Male mice were trained under an FR1 schedule for sucrose, similar to previous experiments with morphine treated mice. Following criteria, mice were given PR tests of motivated behavior 30 minutes after receiving an injection of saline or CNO (2 mg/kg), each session separated 24-h apart.

**Figure 7.**
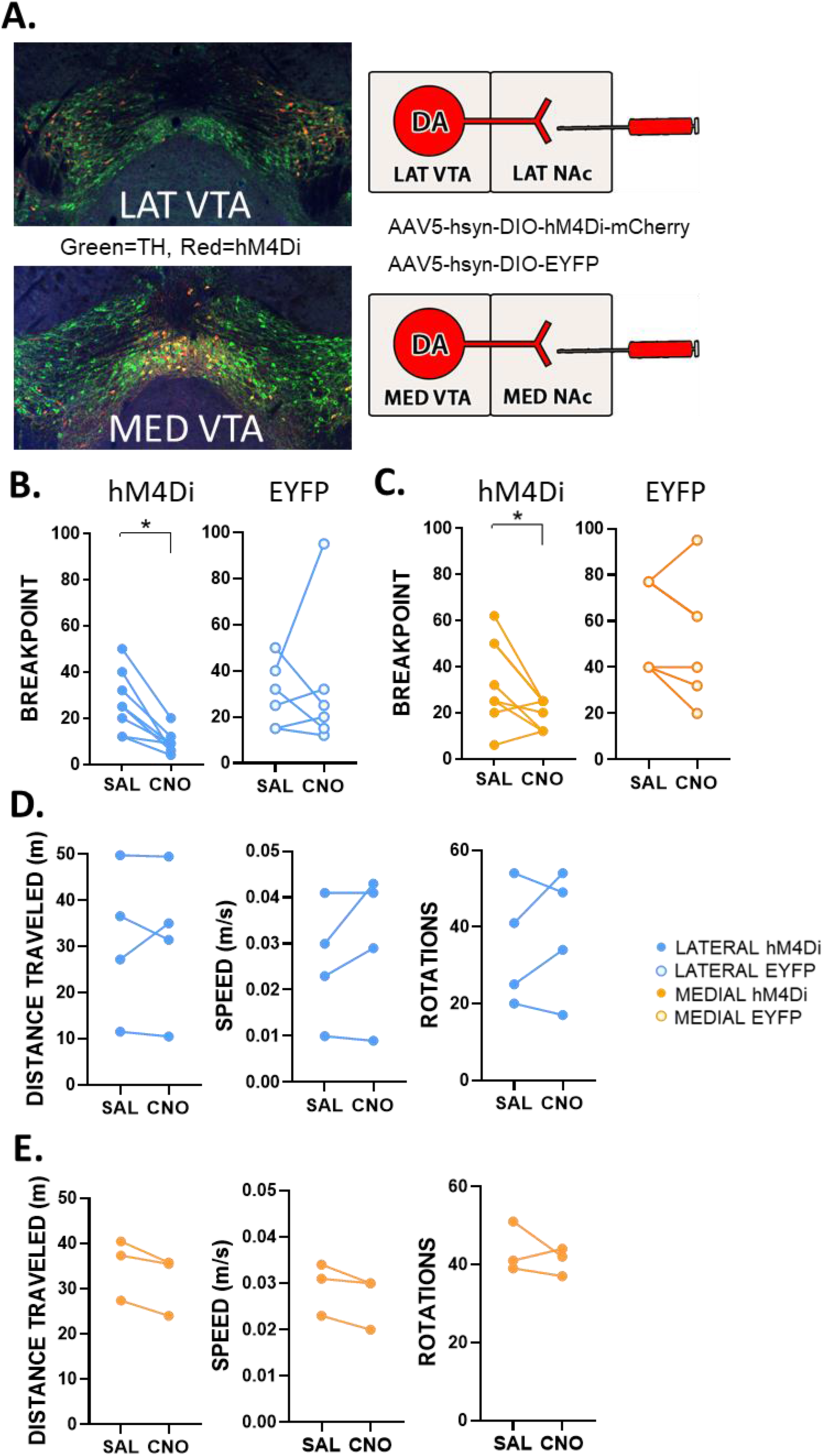
Chemogenetic inhibition of the either mesoaccumbal subcircuit significantly decreases effortful motivation for sucrose. **(A.)** Targeting either the lateral (blue dots) or medial (orange dots) mesoaccumbal subcircuit with Cre-dependent inhibitory DREADDs known for their retrograde transport, sucrose PR tests were run in morphine-naïve DAT-IRES-Cre male mice following both saline and CNO injection (2 mg/kg). Representative images of post hoc histology analysis show double labelled (yellow) cells both positive for tyrosine hydroxylase (green) and hM4Di (red). Doubled labelled cells were found to be biased towards the lateral VTA for lateral NAc shell injections and the medial VTA for medial NAc shell injections. Inhibition of lateral **(B**.) and medial **(C.)** subcircuit DA cells significantly decreased breakpoint values for sucrose reward (N=8 per group), with no effect observed in viral controls in either subcircuit (N=6 per group). Mobility tests showed no appreciable difference between treatment days in distance traveled, speed, or rotations for **(D.)** laterally or **(E.)** medially infused mice. N=3–4 per group. *p< 0.05.

Two-way (subcircuit x virus) RM ANOVA analysis of breakpoint values showed an interaction between day and virus (F(1,24)=4.533, p=.044, η_**p**_^**2**^=.159), and post hoc tests showed a significant decrease in breakpoint values for both medially (F(1,24)=5.431, p=.029, η_**p**_^**2**^=.185) and laterally (F(1,24)=7.616, p=.011, η_**p**_^**2**^=.241) DREADD injected animals across days (**Figure 7B, C**). To ensure decreases in responding were not due to decreases in locomotor activity resulting from chemogenetic manipulations, mobility assessments were taken from a subset of both medially and laterally-targeted mice a few days following progressive ratio testing. Using ANY-maze software to track animal movement recorded from cameras positioned above the animals, speed (m/s), distance traveled (m) and number of rotations were quantified during 20-minute open field tests. Each test followed 30 minutes of either intraperitoneal saline and CNO injections separated by 24-h. Non-parametric paired t-tests (Wilcoxon) were used to calculate significance due to some low sample sizes across viral groups (n≤3). Mobility tests showed no appreciable difference between treatment days in distance traveled (W=-2, p=.8750), speed (W=4, p=.5000), or rotations (W=4, p=.6250) for laterally infused mice (**Figure 7D**). Nor were there differences in distance (W=-6, p=.2500), speed (W=-6, p=.2500) or rotations (W=-2, p=.7500) for medially infused mice (**Figure 7E**).

These data suggest selective inhibition of either mesoaccumbal subcircuit sufficiently decreases motivated behavior for natural reward without affecting mobility.

## DISCUSSION

The results presented demonstrate a profile of sex- and subcircuit-specific synaptic adaptations associated with behavioral changes towards different rewards during protracted morphine withdrawal. While both male and female withdrawn mice increased effortful responding for and consumption of morphine, concurrent decreases in effort towards sucrose were observed only in males. Male exclusive synaptic plasticity to lateral mesaoccumbal subcircuit VTA DA cells was also found to parallel these behavioral changes, resulting in a hypoactive neural state combined with increased inhibitory tone. Overall, this study further characterizes the role DA circuitry has in protracted opioid withdrawal-related alterations in motivated behavior by focusing on treatment effects across sex and mesoaccumbal subcircuit.

Our results showing higher acute somatic withdrawal signs in males agrees with previous data, wherein male rats similarly display elevated symptoms compared to females during the first 48 hours of morphine withdrawal ^41,42^. Such differences in acute withdrawal may be attributed to several sex differences noted in the literature. Relating to pain sensitivity and aversion, males have been shown to have a significantly higher expression of mu opioid receptors (MOR) in the ventrolateral periaqueductal grey (PAG) relative to cycling females, and morphine withdrawal has been associated with the release of proinflammatory cytokines within the spinal cord exclusively in males ^43,44^. Furthermore, MOR gene slice variants, each uniquely correlated to differences in analgesia, itch, opioid tolerance and reward, have been evidenced to be both sex- and brain region-dependent ^42^. Sex differences in regional brain metabolism have also been observed during spontaneous morphine withdrawal, revealing disparate changes in the periaqueductal grey, amygdala, insular cortex, dorsal striatum, and other regions important to various forms of expression of withdrawal ^45^.

The effect morphine withdrawal has on motivated responding for natural reward is contradictorily reported in the literature. Spontaneous morphine withdrawal during both acute (24 hours) and more protracted (10 days) timepoints has been shown to decrease PR breakpoint values for 4% liquid sucrose in rats (Zhang et al., 2007). And 15 days of heroin withdrawal has been shown to decrease breakpoints values obtained for food reward in rats ^47^. Conversely, 8 and 14 days of spontaneous morphine withdrawal in rats has also been shown to have either no effect on motivated responding for sucrose or increase it depending on the sucrose concentration ^27,29^. The cause of this discrepancy is unknown, though slight inconsistencies in experimental conditions such as drug dosage, concentration of sucrose reinforcer, and withdrawal period may all be partially responsible. Both the current study and Zhang et al. show a decrease in post-treatment sucrose breakpoint values in both saline and morphine treated animals. While this treatment-independent decrease is not understood, it is possible that the interim itself between pre-treatment and post-treatment tests – approximately 10-14 days without operant conditioning – influences sensitivity to the PR test. Regardless, the morphine treated animals in both Zhang et al. and the current study displayed more severe reductions to motivated responding than their saline counterparts, indicating a withdrawal-associated effect separate from that related to reinforcement history. Observed decreases in opioidergic signaling following opioid dependence may help explain the found decreases in sucrose motivation. Decreasing endogenous opioids and opioid receptor activation has been shown to decrease both sucrose consumption and motivated behavior in opioid-naïve animals ^48,49^. Chronic opioid exposure has been associated with decreases in opioid receptor expression and availability as well as decreased endogenous opioid peptide gene expression in the VTA ^50–52^. And while heroin IVSA in rodents does not seem to change mu opioid receptor expression in either sex ^53^, protracted heroin withdrawal in male mice results in increased proopiomelanocortin (precursor to endogenous opioid production) cell reporter signal and mRNA levels in the amygdala and hippocampus, respectively ^51^. Such alterations may be compensatory mechanisms against decreased opioidergic signaling. While little is understood about the function of extrahypothalamic proopiomelanocortin expressing neurons and their local peptide production, these data suggest opioid-induced alterations to endogenous opioid signaling within limbic regions may alter reinforcement of palatable food and other affect-related processes.

Results within displayed increases in morphine intake and reinforcement during protracted morphine withdrawal across both sexes. This contrasts with a recent report from Chartoff and colleagues using similar escalated morphine regimens in both sexes, demonstrating a sex-dependent effect on opioid consumption and reinforcement ^26^. Results from Chartoff and colleagues show withdrawn males but not females acquire opioid self-administration faster than their saline counterparts, but no differences in maintenance were observed between treatments groups of either sex. This contrasts with the current study, observing immediate (day 1 of opioid self-administration) higher response rates and infusions for opioids compared to saline controls across sex. Also, while data presented herein display increased breakpoint values obtained by withdrawn animals across sex, the opposite study found a decrease in breakpoints obtained by withdrawn females. These discrepancies may possibly be explained by several experimental differences. In the current study, animals underwent operant conditioning for sucrose before morphine exposure, while Chartoff and colleagues began their experiments with a morphine regimen before operant training, using intravenous opioids to assess acquisition of both opioid intake and the instrumental task itself. Second, while non-contingent morphine was used in both studies to induce dependence, the current study measured IVSA with the same morphine drug while the opposite study used oxycodone. Third, withdrawal timepoints under which measures were taken differed, the current study being 16 days following the last morphine injection, including only 3 days of intravenous morphine self-administration. The contrasting study measured motivated oxycodone self-administration 52 days after the last morphine injection, with 7 weeks of a combination of intermittent and daily access to opioids before motivational tests. It’s also possible that other factors influenced the different results, such as species differences (rats vs mice) and dose of morphine used (the current study used up to 100 mg/kg vs 30 mg/kg used by Mavrikaki et al). Thus, investigation of sex differences in opioid withdrawal behavior may benefit from parsing the effects of operant training using different opioids, different withdrawal timepoints, dose-dependence, and other factors that may impact opioid self-administration.

A motivational shift away from natural rewards and towards morphine was found exclusively in males, suggesting an important opioid-induced sex difference that may confer abuse vulnerability. This runs counter to cumulating evidence outlining females as relatively more susceptible to drug abuse and relapse during opioid withdrawal ^54–56^. However, numerous affective states are believed to impact opioid abuse, each possibly exerting different effects across sexes. For instance, females are believed to be more sensitive to stress- and drug cue-induced opioid relapse compared to males ^57,58^. On the other hand, males may be more prone to pain motives relating to opioid intake, with evidence suggesting they are more sensitive to hyperalgesia following opioid withdrawal and more likely to escalate opioids in response to pain ^43,44,59^. And morphine has been found to induce divergent affective states between sexes, with males reporting greater positive and females greater negative subjective mood-altering effects following acute injection ^60^. It’s also possible that decreases in motivation for natural reward have little impact on opioid reward in animals, and in the case of the current study, did not confer additional drug motive in males. This would be a surprising result, however, as reinforcement of opioids and palatable food are both supported by mesoaccumbal DA ^61^, the system found to be partially suppressed in males during protracted withdrawal. Additionally, neuroadaptations to VTA DA cells have been previously shown to alter behavior towards both natural and opioid reward following endogenous opioidergic signaling, suggesting supranatural levels of opioids such as those used here have the potential to meet if not exceed such effects ^62^. Related alterations in motivated behavior, whereby preference for and consumption of opioids surpasses that of non-drug rewards (e.g., food, exercise), is thought to be a predominant factor conferring vulnerability to escalated drug use and relapse ^28,63–76^. Therefore, additional studies focusing on altered reward learning and reinforcement behaviors between drug and natural rewards is warranted. Dual-choice, mutli-operant paradigms may be fruitful in this regard, allowing choice behaviors during simultaneous self-administration of different reinforcers. For example, male rats undergoing acute fentanyl withdrawal have been shown to display significantly higher choice for fentanyl over food compared to females in a choice paradigm ^77^. Further attention to protracted withdrawal timepoints using similar paradigms seems promising.

Commonly reported symptomatology of opioid withdrawal in clinical studies, especially protracted withdrawal, is negative affect, including depression and anhedonia ^63,75,78,79^. This has been mirrored by preclinical reports of anhedonia-like behavior displayed by animals during several stages of opioid withdrawal ^29,80–83^. Male withdrawn mice decreased their PR breakpoint values for sucrose, but not their preference, in the present study. This suggests a protracted opioid withdrawal-associated deficit in motivation for, but not the hedonic response to, natural sucrose reward. This interpretation both supports and refutes previous findings, as consummatory behaviors measured for withdrawal-related anhedonia in rodents present inconsistent data ^27^. Withdrawal-associated anhedonia alone might not best determine complex constructs like drug intake susceptibility, as, counterintuitively, it has been found to be correlated with a decrease in opioid intake in rats ^84^. Further investigation into the relationship between consummatory, appetitive, and different affective behaviors following opioid withdrawal may help compartmentalize the complex behavioral effects of opioid dependence.

A limitation of the current study lies within potential effects of sugar consumption, as prolonged sucrose intake may confound plasticity effects of opioid dependence on DA cells via altered opioid and DA receptor expression ^85–87^. Sucrose binging has also been found to reliably evoke DA release in the medial NAc shell, and naloxone precipitated withdrawal following prolonged intermittent binging induces both behavioral and DA-depleting effects akin to withdrawal from drugs of abuse ^88,89^. Therefore, potential effects of sucrose intake on the neuroadaptations found in male VTA DA cells may not be ruled out.

A general hypoactive state was displayed in withdrawn male lateral mesoaccumbal subcircuit DA cells, with both a decrease in membrane excitability and an increase in GABA_A_R-dependent mIPSC frequency found. One possible explanation for the increased rheobase is an adapted function and/ or expression of factors supporting intrinsic excitability. DA cell firing properties and responsivity have been shown to rely on several ion channel functions and other postsynaptic mechanisms sensitive to drugs, including opioids ^90–95^. Potassium current alterations may be the most parsimonious candidate, being an opioid-sensitive direct modulator of intrinsic excitability ^96^. Studies of opioid withdrawal-related intrinsic plasticity in VTA DA cells are seriously lacking, however, and will be critical to further characterizing opioid-induced neuroadaptations.

An alternative explanation for the decreased excitability in DA cells seen here may lie within the increased inhibitory tone that accompanies it. While increases in sIPSC frequency were found to be short of significance (p=.061), the selectivity of this trend to male lVTA-lShell DA cells – coinciding with identical male lVTA-lShell selectivity of increased rheobase – suggests it is may be related to protracted morphine withdrawal in males. Blocking GABA_A_ receptors has been found to attenuate increases in rheobase in VTA DA cells during states of high presynaptic inhibition related to pain ^97^. This suggests that increased inhibitory tone in the VTA may decrease DA cell excitability, which could feasibly be the case for the current study and may explain the coincidence of both synaptic effects related to protracted withdrawal. Additional experiments controlling for GABA_A_ current during measures of excitability are warranted in this context. If withdrawal-associated increases in rheobase do not depend on inhibitory tone, it may then be possible to separate differences in presynaptic (inhibition) and postsynaptic (intrinsic excitability) effects. However, if increases in rheobase are a result of increased inhibitory tone, this may amount to a simplification of mechanistic and therapeutic targets in the context of opioid withdrawal.

Interpretation of the subcircuit- and sex-specific neuroadaptations observed in the current study should be limited to a protracted withdrawal profile. During acute opioid administration, for instance, the medial but not lateral mesoaccumbal subcircuit has been demonstrated to mediate the reinforcing effects of heroin in mice. Lüscher and colleagues showed that opioids disinhibit the medial mesoaccumbal subcircuit via MOR activation on VTA GABA afferents, in turn stimulating DA release in the medial shell to drive behavioral output ^15^. This conclusion contrasts with the current lateral subcircuit-specific plasticty effects and raises questions surrounding period or timepoint of drug use. For instance, if the medial subcircuit is initially engaged by the robust effect of opioids, does it undergo its own drug-induced neuroadaptations, and if so, when? Also, at what timepoint during withdrawal (acute vs protracted) does increased inhibition in the lateral subcircuit begin? Such questions draw attention to the time-dependent nature of opioid withdrawal-associated plasticity changes in the VTA – an understudied but important characteristic that may be key to understanding OUD ^36,98^.

The current study found both the lateral and medial mesoaccumbal DA subcircuit are necessary for intact motivated responding for sucrose in males. This agrees with previous reports describing motivated behavior as a composite function driven by different DA cell subpopulations, and as a result shedding light on the role of lVTA-lShell DA neurons in reinforced behavior ^6,39,99,100^. As subcircuit-specific chemogenetic manipulations were performed only in males, and as very little sex-dependent mesoaccumbal subcircuit analysis has been reported, it’s yet unknown if either lVTA-lShell or mVTA-mShell DA cells diverge in their ability to reinforce behavior across sex. This would not be surprising considering sexually divergent basal properties in VTA DA cells have been described, albeit without employing subcircuit analysis ^101,102^. In addition, sex-specific neuroadaptations following protracted remifentanil withdrawal have been observed, for instance, within pyramidal neurons of the prefrontal cortex ^23^. Sex differences were found between treatment groups in several neuroadaptations that depended on drug schedule (10-16 days vs 25-30 days IVSA) as well as the specific type of measure taken (i.e., membrane excitability vs somatodendritic current).

Regarding mesoaccumbal circuitry, a sex- and time-dependent adaptation to VTA DA cell activity has been reported by Jones and colleagues, with opioid-evoked phasic activity of DA cells bidirectionally changing across a three-week period of opioid exposure in males ^59^. Additionally, Jones and colleagues have shown that long-access heroin self-administration enhances phasic DA release in the medial NAc shell but not core subregion in females, with males displaying no difference in either subregion ^53^. Despite these recent informative investigations of selective opioid effects, there has been very little investigation of sex-dependent VTA DA plasticity mechanisms following opioid exposure, even less following protracted withdrawal. Thus, future investigations of time- and sex-dependent neuroplasticity events in the mesoaccumbal subcircuits is highly warranted. The male-exclusive behavioral effects observed during protracted withdrawal emphasize the importance of the diversity of biological factors impinging on opioid-induced effects (sex, withdrawal phase, signaling system, subcircuit plasticity, etc.,). Isolating biological characteristics of withdrawal specific to such factors may yield effective therapeutic targets against neurobehavioral maladaptations associated with opioid withdrawal and OUD.

## Supplemental Material

**Figure S1.**
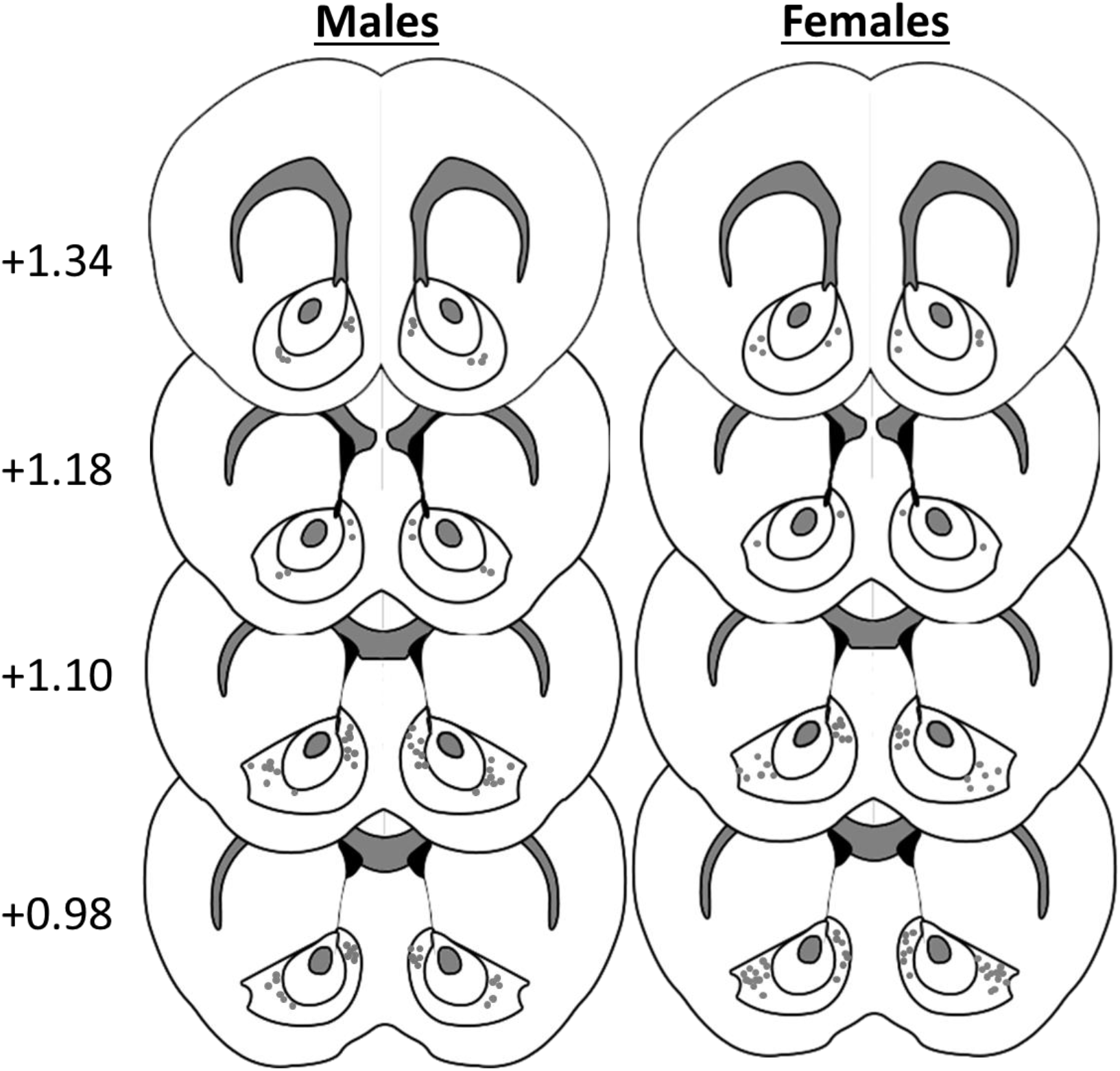
Injection placements of Retrobead™ injections within the NAc prior to electrophysiological recordings. All coordinates referenced with respect to Bregma (mm).

**Figure S2.**
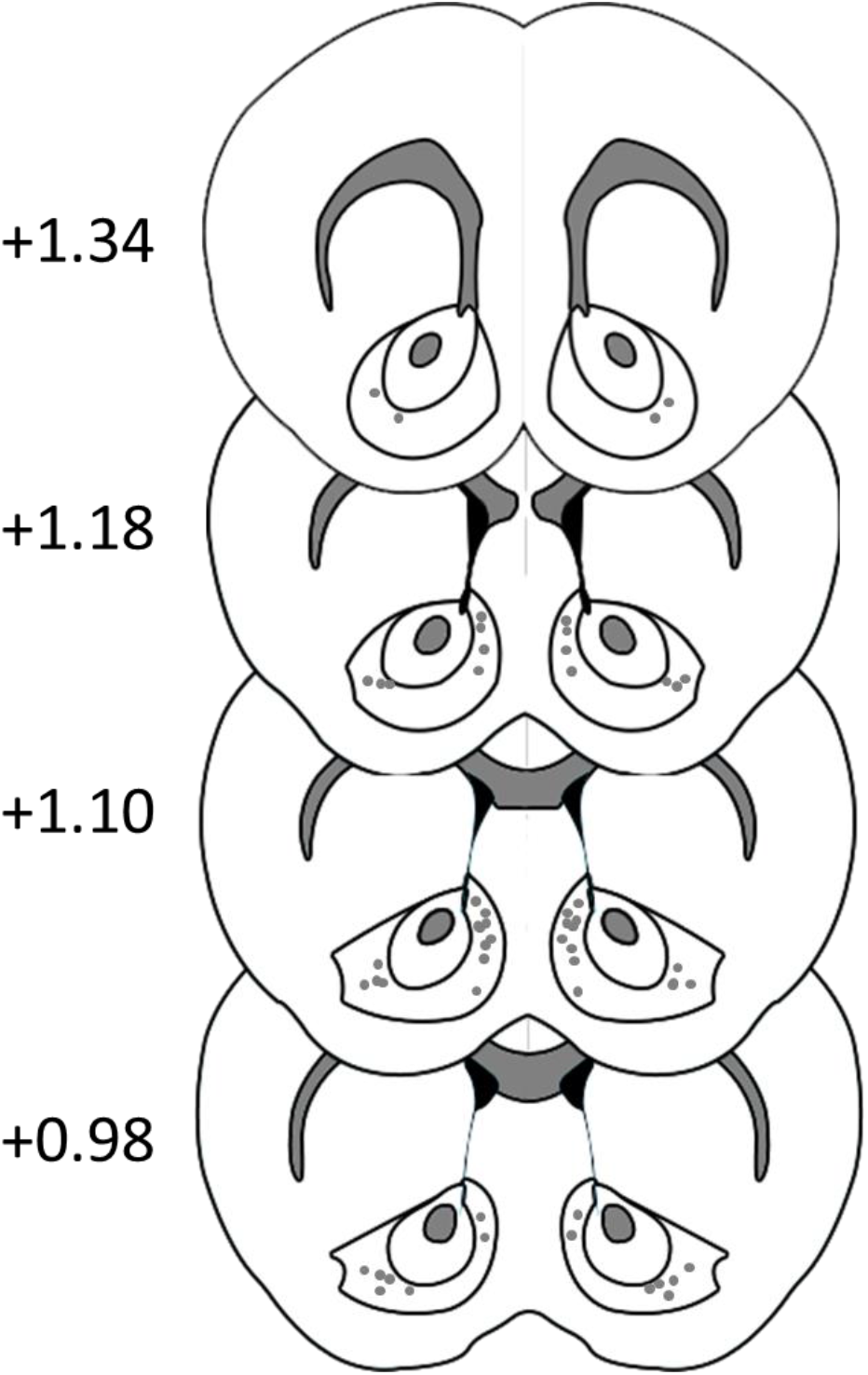
Injection placements of hM4Di & EYFP injections within the NAc prior to chemogenetic studies. All coordinates referenced with respect to Bregma (mm).

